# The barley ear row-number allele ancestral to the six-row allele *vrs1.a1* is found in wild barley from the Fertile Crescent

**DOI:** 10.1101/2025.11.20.688930

**Authors:** James A. Bedford, Huw Jones, the AEGIS Consortium, James Cockram

**Affiliations:** Niab, Park Farm, Villa Road, Histon, Cambridge, CB24 9NZ, United Kingdom

**Keywords:** Barley genetic diversity, crop domestication, haplotype network, single nucleotide polymorphism (SNP), *Six-rowed spike 1* (VRS1), yield components

## Abstract

Barley (*Hordeum vulgare* L.) exhibits two main inflorescence phenotypes: the wild-type ‘two-row’ form in which only the central spikelet at each rachis node is fertile, and the ‘six-row’ form, where mutations in the homeobox gene *Six-rowed spike 1* (*VRS1*) confer fertility to all three spikelets. While the six-row alleles *vrs1.a2* and *vrs1.a3* arose independently from point mutations in the ancestral *Vrs1.b2* and *Vrs1.b3* alleles, the origin of the hypothesised wild-type *Vrs1.b1* allele ancestral to *vrs1.a1* is unclear. To explore the origin and exploitation of *VRS1* alleles, we re-sequenced *VRS1* in 98 cultivars, 170 landraces and 69 wild barley accessions, identifying 39 haplotypes. Sequence analysis confirmed *vrs1.a1* as the most commonly used six-row allele in European cultivars. Subsequent analysis of the landrace and wild barley data identified three occurrences of the haplotype consistent with a *vrs1.b1* allele, all from wild barley. These wild barley accessions originated from the eastern Fertile Crescent (Iran) and the Caspian Sea region, suggesting *vrs1.a1* arose via mutation of *Vrs1.b1* within barley’s domestication centre - unlike *vrs1.a2* and *vrs1.a3*, which originated in the Western Mediterranean and East Asia cultivated barley genepools, respectively. In addition, we identified ten novel VRS1 amino acid changes in wild barley accessions, which given the pleiotropic effects of *VRS1* on traits such as leaf size, vein number and tiller number, may be of interest for future functional investigation. Overall, this study provides insights into the evolution, domestication and utilisation of genetic variation at *VRS1*, a key gene influencing barley architecture and agricultural performance.

## Introduction

Barley (*Hordeum vulgare* L.) is a globally important cereal crop with a worldwide annual production of ∼146 million tons, and is the world’s fourth most important cereal crop after maize, wheat and rice (FAO, 2024). Barley cultivars can broadly be divided into two main groups, based on their end-use. Two-row barley, in which each rachis node of the inflorescence contains a central fertile floret flanked by two sterile florets, is predominantly used for brewing (Figure 1). In contrast, six-row barley, in which all three florets at each rachis node are fertile, is mostly used for animal feed (Figure 1). The wild progenitor of barley (*H. vulgare* ssp. *spontaneum* K.Koch) has the two-rowed ear phenotype (Figure 1), and is native to the Eastern Mediterranean, Central Asia, the Tibetan Plateau and China. Work in cultivated barley has shown that mutation at the homeodomain-leucine zipper I-class homeobox gene *Six-rowed spike 1* (*VRS1*) results in conversion of the wild-type two-row phenotype to the six-row form (Komatsuda et al. 2007). Three mutant alleles have been described that result in a six-row phenotype (Bregitzer et al. 2007), each of which results in disruption of the VRS1 predicted protein (Komatsuda et al. 2007). In the *vrs1.a1* allele, a single nucleotide deletion at position 681 (*G681/Del*) relative to the wild-type allele from cv. ‘Bonus’ leads to a frameshift within the recognition helix and premature termination of the predicted protein. The *vrs1.a2* allele carries a 1 bp insertion within exon-2 (*T243/Ins*) that leads to a frameshift after amino acid 40 and premature truncation of the predicted protein before the start of the homeodomain. Finally, the *vrs1.a3* allele has a missense mutation (*C349/G*) in exon-2 that results in an amino acid substitution at a highly conserved amino acid residue within the homeobox domain (F75/L).

**Figure 1.**
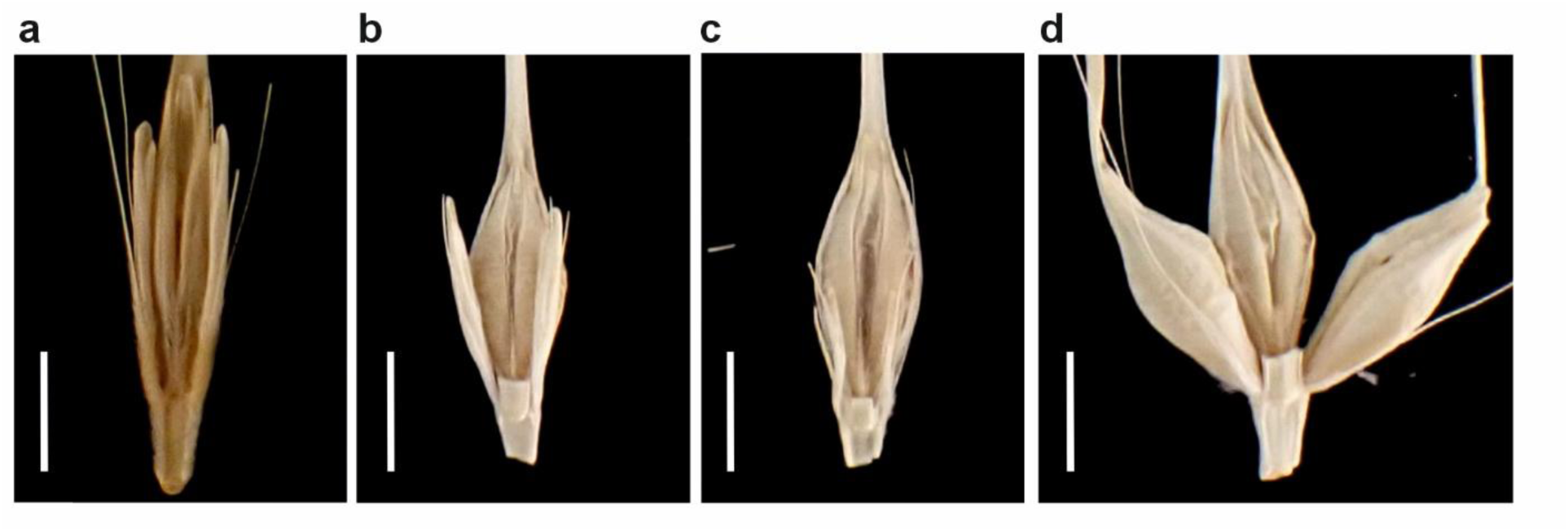
Examples of different barley spikelet morphologies controlled by *VRS1*. Scale bar = 5 mm. (a) Wild barley (*Hordeum vulgare* ssp. *spontaneum*) two-row form (HorID accession 10924. *Vrs1.b2* allele) in which at each rachis node, the central fertile floret is flanked by two sterile florets. (b) The two-row type in barley cv. ‘Maris Otter’ (allele *Vrs1.b3*). The *deficiens* type in cv. ‘Craft’ (*Vrs1.t1* allele) in which the two sterile florets flanking the central fertile floret are severely reduced in form. (d) The six-row type in cv. ‘KWS Feeris’ in which all three florets per rachis node are fertile.

Barley is thought to have been domesticated from its wild progenitor in the Neolithic period in the Fertile Crescent region of the Middle East (Fuller et al. 2023; Guo et al. 2025; Hillman, 1975; Zohary et al. 2012). Post-domestication, human activity lead to the spread of barley cultivation from its origins in the Fertile Crescent, first into Europe, Asia and Northern Africa (Zohary et al. 2012), and subsequently to the Americas and Oceania. While the first domesticated barley (∼10,000 years before present) was two-rowed, around 1,000 years later six-row types had begun and soon largely replaced the two-row cultivated form, becoming the most important crop in the Near East during the Neolithic (Zohary et al. 2012; Harlan, 1968; Helbaek, 1959; Pourkheirandish and Komatsuda, 2007). Prior to the use of crop cultivars with advent of industrial plant breeding approaches in the late 19^th^ Century, the post-domestication spread of barley was via use and transport of germplasm stocks now termed ‘landraces’ (germplasm grown and selected by growers for use in their local agro-ecological regions of cultivation and for specific end-uses) (Jones et al. 2011). Analysis of genetic markers and *VRS1* haplotypes associated with the different *Vrs1* alleles in barley cultivars, landraces and wild barley has found that the six-row alleles *vrs1.a2* and *vrs1.a3* likely arose post-domestication in cultivated barley via single mutations in their respective wild-type alleles (Komatsuda et al. 2007; Saisho et al. 2009; Tanno et al. 2002). Specifically, *vrs1.a2* is thought to have directly arisen from *Vrs1.b2* in barley cultivated in the Western Mediterranean, and *vrs1.a3* from *Vrs1.b3* in East Asia. However, while *vrs1.a1* is the most commonly used six-row allele in modern barley cultivars (Casas et al. 2018; Komatsuda et al. 2007; Saisho et al. 2009), the hypothetical progenitor *Vrs1.b1* allele has not been identified in cultivated barley.

Allelic variation at *VRS1* also affects additional barley traits. For example, the extreme suppression of the infertile lateral spikelets observed in the naturally occurring *deficiens* allele *Vrs1.t1* (Figure 1c), which carries three amino acid substitutions in its predicted protein compared to the wild-type allele from cv. ‘Bonus’ (D8/G, D26/E and S184/G), has been shown via comparison to the induced *deficiens* mutants *Def2* (*Vrs1.t2*, S184/G) and *Def3* (*Vrs1.t3*, S186/P) to be due to the *S184/G* missense mutation at a conserved amino acid position towards the C-terminus of the VRS1 protein (Sakuma et al. 2017). The six-row *vrs1.a1* allele also confers increased leaf size, predominantly due to an increase in leaf width, as well as increased leaf vein number and percentage nitrogen content (Thirulogachandar et al. 2017). *VRS1* is expressed during leaf primordia initiation and development where it represses the proliferation of leaf and lateral spikelet primordia (Sakuma et al. 2017; Thirulogachandar et al. 2017). The six-row allele *vrs1.a1*, as well as the intermedium alleles *vrs1*(*int-d.11*) and *vrs1*(*int-d.22*), are also known to confer reduced tiller number at maturity (Liller et al. 2015; Thirulogachandar et al. 2017). This reduction in tiller number is not observed at earlier stages of development, indicating *VRS1* does not affect the initiation and initial growth of tillers. The increased grain number per spike and reduced absolute spikelet abortion observed in six-row types, despite the presence of three times the number of fertile spikelets compared to two-row barley, is hypothesised to be due, at least in part, by the increased leaf area found I n six-row types (Alqudah and Schnurbusch, 2015; Thirulogachandar et al. 2017). Given the numerous positive effects of six-row *vrs1* alleles on ear and leaf traits, the reduction of tiller number at later stages in the plant lifecycle are thought to be due to negative trade-off interactions that occur after tiller initiation (Thirulogachandar et al. 2017).

Given the key role of *VRS1* in the barley domestication and post-domestication process, and the emerging findings that *VRS1* has pleiotropic effects on other traits of potential agronomic interest in barley, here we screen a panel of wild, landrace and cultivated barley accessions previously phenotyped for ear-row type with the aim of (i) Identifying sources of VRS1 missense variation in the wider barley genepool, and (ii) resolving the evolutionary origin of the six-row *vrs1.a1* allele.

## Materials and Methods

### Barley germplasm and DNA extraction

Barley scientific name nomenclature follows that outlined by POWO (2025). Seed for 69 wild barley and 170 barley landrace accessions was obtained from the genebanks listed in Supplementary Table S1, as previously described by Jones et al. (2008; 2011). Additionally, seed for 98 of the barley cultivars described by Rostoks et al. (2006) were sourced from the James Hutton Institute, Dundee, UK (Supplementary Table S1). Genomic DNA was extracted from seedling leaf material collected from a single plant per accession using a DNAeasy Kit (Qiagen), following the manufacturer’s instructions. DNA concentration was estimated using a NanoDrop spectrophotometer (Thermo Fisher Scientific) and diluted to 10 ng/μl using molecular biology-grade water (Thermo Fischer Scientific).

### VRS1 PCR amplification and sequencing

*VRS1* was amplified via polymerase chain reaction (PCR) using the primers OSU-VRS1-F1 (5’-CCGATCACCTTCACATCTCC-3’) and OSU-VRS1-R1 (5’-GGTTTCTGCCGATCTTGAAGC-3’). PCR amplification was undertaken in 10 µl reactions using the FastStart *Taq* DNA polymerase kit (Roche) and a a Veriti 96-well Thermo Cycler (Applied Biosystems) using the following PCR cycle: 6 min at 96 °C, followed by 35 cycles of 50 sec at 96 °C, 50 sec at 60 °C, 90 sec at 72 °C, and a final extension of 7 min at 72 °C. PCR products were subsampled to check for adequate amplification and expected product size via electrophoresis across ethidium bromide stained 1% agarose gels alongside a 1 kb DNA ladder. Amplification products were ExoSap treated in reactions containing 1.5 U of Exonuclease I (New England Biolabs) and 1 U of Shrimp Alkaline Phosphatase (Promega) and incubated at 37 °C for 60 min followed by 15 min at 96 °C using a Veriti 96-well Thermo Cycler. Sanger sequencing was then undertaken using the ExoSap treated PCR products using BigDye Kit v3.1 (Applied Biosystems) in 10 μl reactions following the manufacturers’ instructions, and the resulting fluorescently end-labelled single stranded extension products separated and visualized using a 3730 DNA Analyzer (Applied Bio-systems). Five primers were used for Sanger sequencing: OSU-VRS1-F1, OSU-VRS1-R1, HvVRS1-F1 (5’-CAGAACAACCTACCGTGTCT-3’), HvVRS1-F2 (5’-AGATGGACGGAGGAGGGGAC-3’) and HvVRS1-R1 (5’-TGTCATCAGCTTAGCCAGCA-3’). DNA sequence traces were manually quality controlled and contigged using Contig Express (Thermo Fischer Scientific). *VRS1* DNA sequence polymorphisms, including single nucleotide polymorphisms (SNPs) and 1-2 bp insertion/deletions (InDels), were positioned with reference to the genomic sequence of the two-row *Vrs1.b3* allele from barley cultivar ‘Bonus’ (GenBank accession AB489121), with the A of the ATG start codon representing position 1 bp.

### Bioinformatic, haplotype and geographical analyses

*VRS1* DNA sequences, and their corresponding predicted protein sequences determined using EMBOSS Transeq, were aligned using Clustal Omega (Madeira et al. 2024). Genomic *VRS1* sequences were used for haplotype analysis using the software PopArt, which implements the minimum spanning tree method (Leigh and Bryant, 2015). Using the collection location latitude and longitude information Supplemental Table S1), the *VRS1* haplotypes for the wild and landrace barley accessions were plotted on geographic maps using the packages ggplot2 (Wickham, 2016) and rnaturalearth (Massicotte and South, 2025) in R version 4.4.0 (R Core Team, 2024). Visualisation of *VRS1* haplotypes across germplasm groups was conducted using the R packages ggplot2 and ggalluvial (Brunson and Quentin, 2023). Two-row wild-type (*Vrs1.b2*, *Vrs1.b3*), six-row mutant (*vrs1.a1, vrs1.b1, vrs1.c1*) and two-row *deficiens* (*Vrs1.t1*) alleles are designated as previously described (e.g. Komatsuda et al. 2007); *VRS1* haplotypes are designated here based on the allele nomenclature in non-italic, along with the inclusion of a numerical suffix (.X, where X = a number). Within each major haplotype (e.g. Vrs1.a1), sub-haplotype numbering is sequential based on position of the first DNA variant in the sequenced region that differs from the reference allele used here for variant calling (e.g. Vrs1.a1.1, for which the first DNA variant which distinguishes it from the canonical Vrs1.a1 haplotype is A-189/G, located in the *VRS1* promotor region). Protein 3D structure was investigated using the AlphaFold protein models for *VRS1* gene model *HORVU.GOLDEN_PROMISE.PROJ.2HG00164530.1* from two-row barley cv. ‘Golden Promise’ (genome assembly: Hvulgare_cv_Golden_Promise_BPGv2, INSDC Assembly, European Nucleotide Archive assembly GCA_949783185.1), accessed using Ensembl Plants (Yates et al. 2022).

## Results

### VRS1 haplotype analysis in cultivated, landrace and wild barley

To provide base-line data from the modern cultivated barley genepool, sequencing *VRS1* in the 98 predominantly European cultivars from the ‘pre-AGOEB’ barley collection was undertaken, with haplotype analysis on the resulting nine DNA variants identified (Figure 2a) finding all to possess one of six common *VRS1* alleles (Figure 2b-c; Supplementary Table S1-S2). Amongst the 82 two-row cultivars analysed, four were phenotypically *deficiens* and carried a *VRS1* haplotype consistent with the *Vrs1.t1* deficiens allele. Of the remaining two-row cultivars, one carried the *Vrs1.b2* haplotype (cv. ‘Camargue’, a German spring barley variety released in 1986), while the remaining 78 cultivars all carried the *Vrs1.b3* allele. For the 16 six-row cultivars, *VRS1* haplotype analysis found roughly equal numbers of cultivars to carry the six-row alleles *vrs1.a1* (five cultivars), *vrs1.a2* (five cultivars) and *vrs1.a3* (five cultivars and 1 morphological stock).

**Figure 2.**
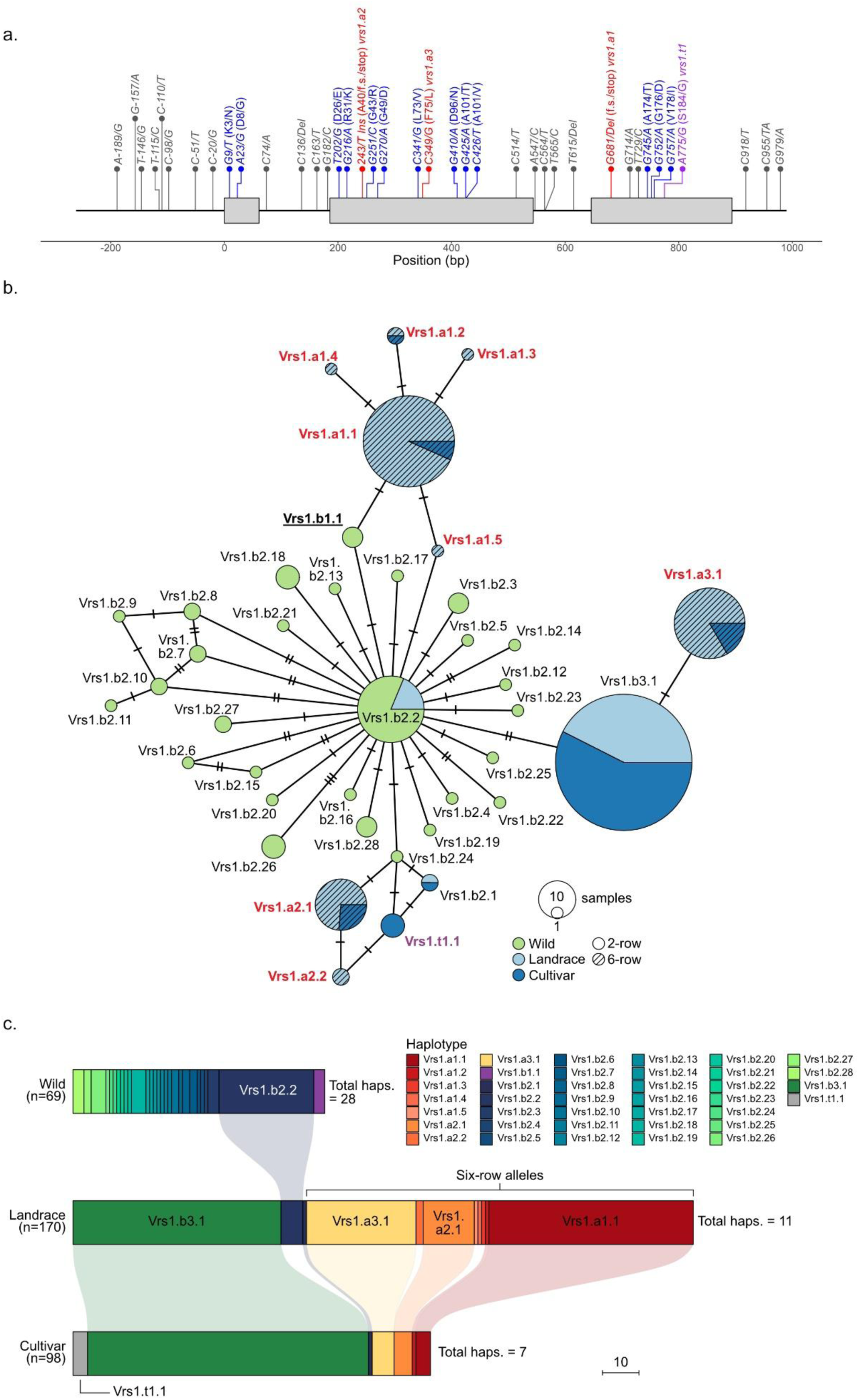
*VRS1* DNA variants and haplotypes identified in wild, landrace and cultivated barley accessions. (a) Positions of *VRS1* DNA variants, relative to the *Vrs1.b3* two-row allele from cv. ‘Bonus’ (GenBank accession AB489121). Variants leading to an amino acid substitution are indicated in blue, those resulting in the six-row phenotype in red and the *Vrs1.t1* deficiens allele highlighted in purple. (b) *VRS1* haplotype network indicating the relationships between the 39 haplotypes identified. The number of DNA variants distinguishing linked haplotypes is indicated using the short bisecting lines. (c) *VRS1* haplotype proportions across the three barley genepools.

Next, *VRS1* sequencing in a collection of 170 European barley landraces (identifying 11 DNA variants) and 69 wild barley accessions from across its natural range (33 DNA variants; Figure 2a) allowed haplotypes and corresponding alleles to be determined across all 239 accessions (Figure 2b; Supplementary Table S1-S2). Similar to the finding in modern varieties, the most common two-row landrace allele as determined by *VRS1* haplotype was *Vrs1.b3* (57 accessions, representing 90% of the two-row barley landraces), with the remaining carrying either *vrs1.b2* (HorID 1393 from France) or a sub-haplotype of the *vrs1.b2* allele which lacks the intron-2 *C564/T* SNP (six accessions, from Syria Afghanistan, Czech Republic ad Bolivia) and termed here haplotype Vrs1.b2.2 (Figure 2b-c). Of the 107 six-row landraces, 53% carried haplotypes consistent with a *vrs1.a1* allele (57 accessions), while the remaining possessed one of the following four sub-haplotypes: Vrs1.a1.2 (HorID 2672, carrying an *A-189/G* SNP in the upstream region), Vrs1.a1.3 (HorID 47, carrying a *C-20/G* SNP in the upstream region), Vrs1.a1.4 (HorID 1173 from Ukraine, the only landrace to carrying a missense mutation: SNP *C426/T*, amino acid substitution A101/V), and Vrs1.a1.5 (HorID 1214, which lacked the 3’UTR *C918/T* variant found in all other Vrs1.a1 haplotypes) (Figure 2b). In the landraces, haplotype analysis identified just one accession predicted to carry the two-row *Vrs1.b2* allele (HorID 1393. Collection location: France), while analysis of haplotype genealogy found no landraces carried a *VRS1* haplotype directly ancestral to the six-row *vrs1.a1* allele (Figure 2b).

*VRS1* haplotype analysis found 66 of the 69 two-row wild barley accessions to possess haplotypes indicative of the wild-type *Vrs1.b2* allele (Figure 2b; Supplementary Table S1-S2). A total of 26 Vrs1.b2 sub-haplotypes were identified, which possessed between 1-3 DNA variants relative to the Vrs1.b2.1 haplotype found in cultivated barley. Twenty-seven of the 28 wild barley Vrs1.b2 sub-haplotypes lacked the intron-2 *C564/T* SNP found in cultivated barley. The exception was a wild barley accession from Cyprus (HorID 4042), which although it possessed the intron-2 *C564/T* SNP, also contained a 3’ UTR SNP (*G997/T*) that differentiated it from the canonical Vrs1.b2.1 haplotype found in modern barley. The most common wild barley haplotype was Vrs1.b1.2, which was present in 30% of the accessions screened, with the remaining Vrs1.b1 sub-haplotypes representing rare occurrences, each present in between 1-4 accessions. Notably, analysis of *VRS1* haplotype genealogy identified three wild barley accessions as carrying a haplotype consistent with the hypothesised *Vrs1.b1* allele directly ancestral to the six-row allele *vrs1.a1* (the most common mutant six-row allele found in cultivated barley) (Figure 2b). All three of the wild barley accessions carrying the Vrs1.a1 haplotype originated within the Fertile Crescent: two from Iran and one from Turkmenistan (HorIDs 3852, 4054 and 3948, respectively) (Figure 3).

**Figure 3.**
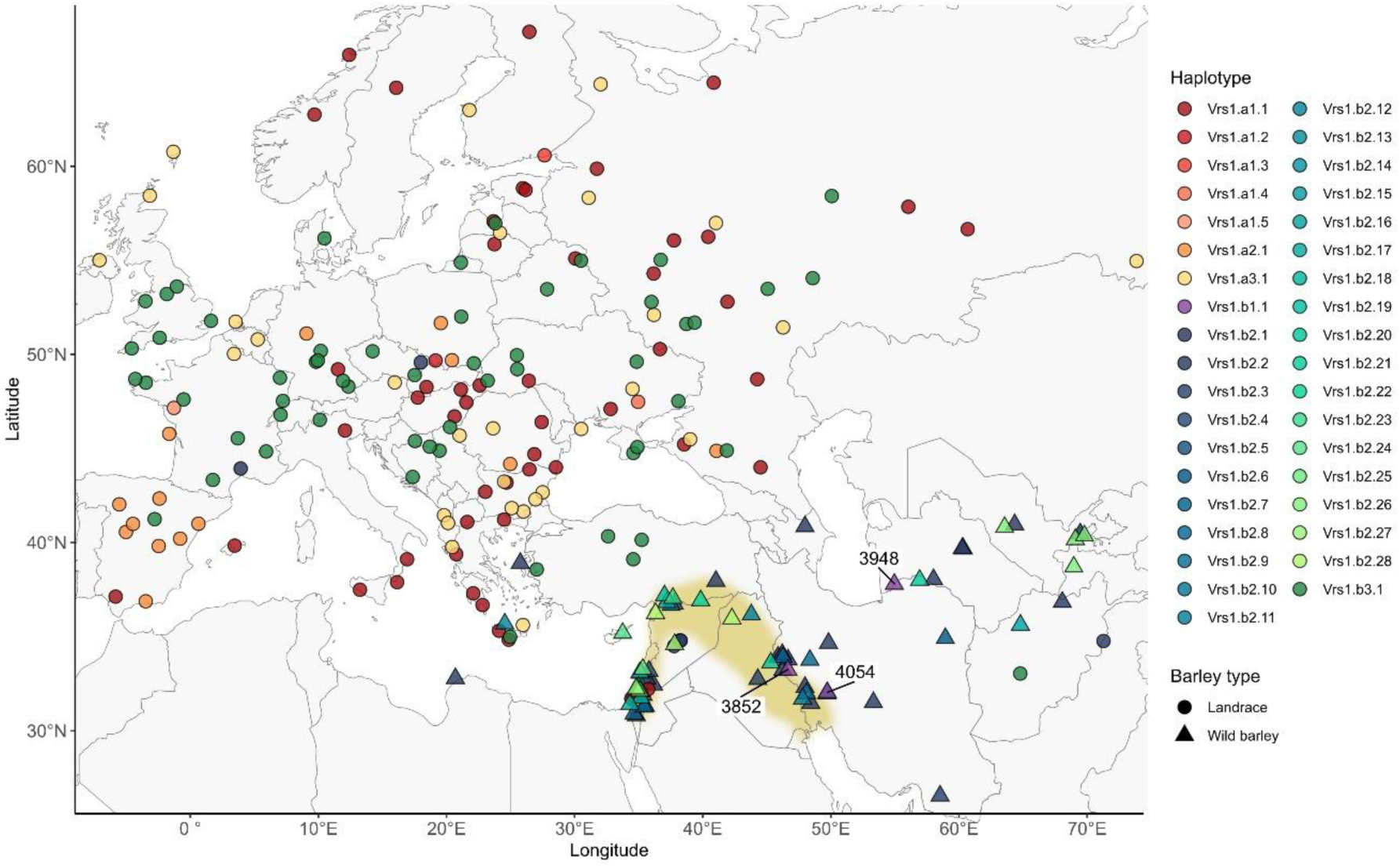
Geographic locations of the landrace (*n* = 161) and wild barley (*n* = 69) accessions with collection location information, indicating their *VRS1* haplotypes. Haplotypes corresponding to the wild-type *Vrs1.b1*, *Vrs1.b2* and *Vrs1.b3* two-row alleles are indicated using the blue-to-green colour scale. Haplotypes corresponding to the mutant *vrs1.a1*, *vrs1.a2* and *vrs1.a3* six-row alleles are indicated using the red-to-yellow colour scale. The three wild barley accessions (HorID 3852, 3948, and 4054) found to carry the Vrs1.b1.1 haplotype ancestral to the mutant *vrs1.a1* six-row allele are indicated. The region representing the Fertile Crescent is shaded in brown.

### Landrace and wild barley VRS1 missense mutations

Overall, 42 SNPs or InDels across the 1,251 bp sequenced region of *VRS1* were identified in the landrace and wild barley accessions. *VRS1* haplotype diversity decreased between the wild barley and landrace genepools (27 and 11 haplotypes, respectively), with a further contraction in diversity found across the transition from landrace to modern cultivars (7 haplotypes) (Figure 2c). While much novel variation was identified at the DNA level in landrace and wild barley accessions, only a subset of variants resulted in novel changes to the predicted VRS1 protein. While these haplotypes conferred wild-type two row alleles, the increasing evidence of pleiotropic effects of *VRS1* on traits other than row number means that such mutations may be of wider interest. In wild barley, all wild-type two-row *VRS1* haplotypes contained the two missense mutations that lead to amino acid substitutions D8/G and D26/E which in modern two-row barley cultivars are only found in the relatively rare *Vrs1.b2* allele (Figure 2b-c, Table 1, Supplementary Table S1). In addition to these, 10 two-row Vrs1.b2 sub-haplotypes contained missense mutations that resulted in a change in amino acid residue elsewhere within the predicted protein. Of these, six mutations were located either within the within the N-terminus (K3/N, R31/K, G43/R) or C-terminus (A174/T, G176/D, V178/I) regions. While both have low model confidence scores in the VRS1 AlphaFold 3D protein model (pLDDT ≤ 52), the acid substitutions in the C-terminus region are located at residues with good levels of amino acid conservation across plant VRS1 homologues (Figure 4). In contrast to the unstructured N- and C-terminus regions, the predicted homeobox-leucine zipper domain (Panther ID PTHR24326) has high AlphaFold 3D model confidence (pLDDT typically > 90 across almost all but the first 15 and last six amino acid residues of the domain). This domain is predicted to consist of three principal structures: the first and second alpha-helixes, which serve as primary structural elements, and the recognition helix which determines DNA binding and specificity. Four *VRS1* haplotypes were found to contain amino acid changes within the homeobox-leucine zipper domain at amino acid residue with high conservation in plant VRS1 homologues: missense mutation L73/V (haplotype Vrs1.b2.15, 68% conservation) located towards the end of the first alpha-helix, D96/N (haplotype Vrs1.b2.20, 100% conservation) located in the connecting loop between the second alpha-helix and the recognition helix, and A101/T (haplotype Vrs1.b2.21, 100% conservation) close to the start of the recognition helix itself (Figure 4; Supplementary Figure S1).

**Figure 4.**
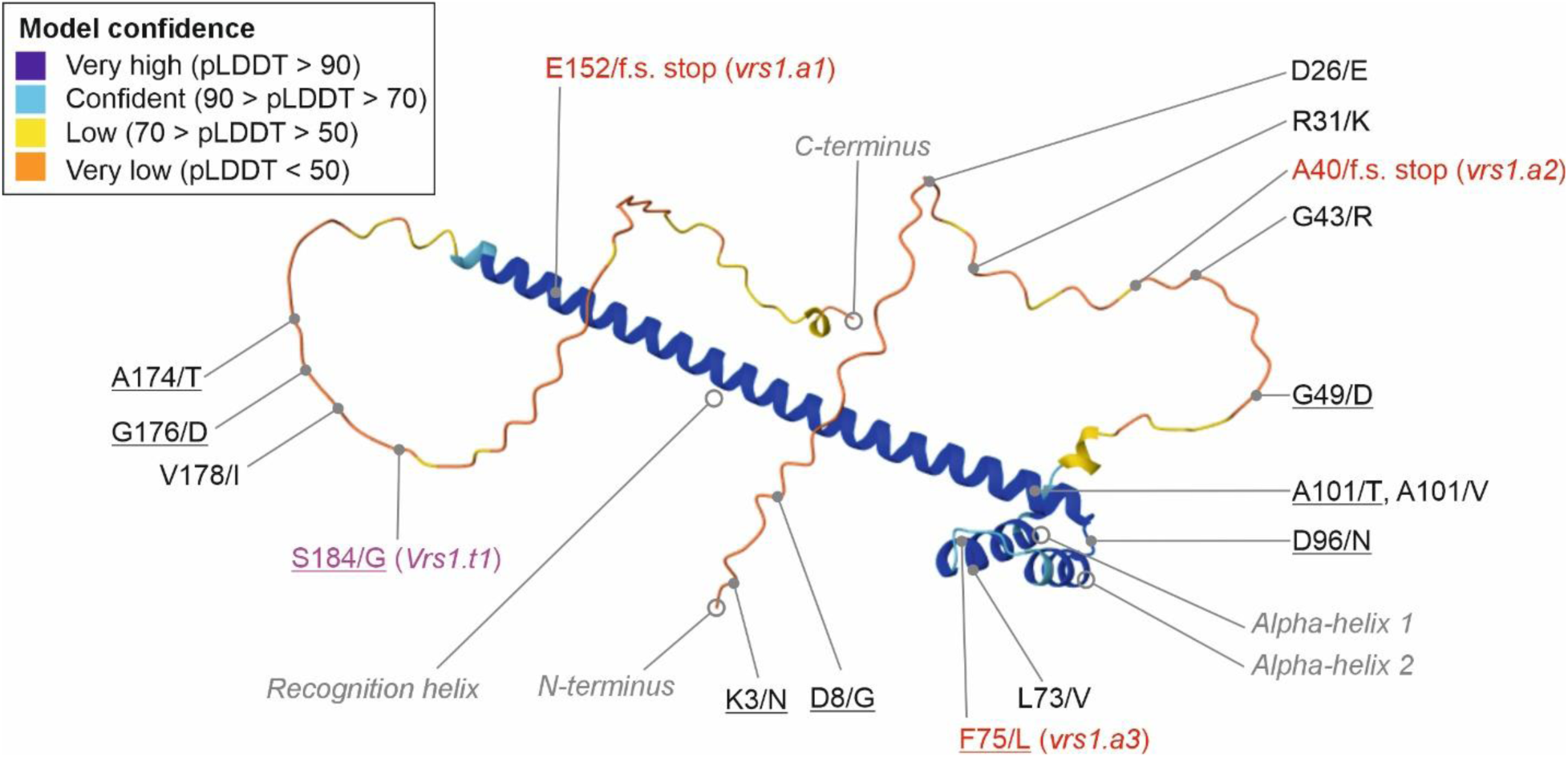
VRS1 AlphaFold 3D protein model for two-row barley cultivar ‘Golden Promise’ (UniProt accession D2KV17, encoded by the *Vrs1.b3* allele) overlaid with the positions of the amino acid substitutions coded for by DNA variants identified in the wild, landrace and cultivar barley genepools investigated. 3D protein model confidence is colour coded as indicated in the key. The homoeobox-leucine zipper domain (Panther ID PTHR24326) overlaps with the ‘very high’ protein model confidence, and consists of alpha-helix 1, alpha-helix 2, and the recognition helix. Non-conservative substitutions are underlined. Mutations causative for the three known six-row alleles are indicated in red. The mutations underlying the two-row *deficiens* allele is indicated in purple. f.s. = frame shift. Stop = premature stop codon.

## Discussion

Barley domestication began in the Near East around 10,000 years ago with the shift from hunter-gatherer societies to farming (Zohary et al. 2012). Rather than a single event, crop domestication is a process that continues to the present day. Key points across this continuum include (i) Neolithic selection for non-brittle rachis types controlled by the *Non-brittle rachis 1* (*btr1*) and *btr2* loci (Pourkheirandish et al. 2015), (ii) the rise of industrial breeding in the late 19^th^ Century (Holmes, 2025), (iii) yield gains from reduced plant stature during the Green Revolution (Hedden, 2023), and (iv) the optimisation of modern-day breeding approaches that deliver a ∼1% yield genetic gains annually (Mackay et al. 2011). Two-row barley first appears in the archaeobotanical record around 10,000 years ago from sites in Syria (van Zeist and Bakker-Heeres, 1985) and Iraq (Helbaek, 1959; Braidwood, 1960), with six-row mutants emerging as key Neolithic innovations in the near east from around 9,000 years ago onwards (Helbaek, 1964; Fairbairn et al. 2002, 2007). Six-row barley is thought to have largely replaced two-rowed types as cultivation first spread beyond the Fertile Crescent (Costantini, 2006; Helbaek, 1959). Today, six-row barley is cultivated in temperate regions across the world, and is used predominantly for animal feed. Accordingly, sequence variation at *VRS1* has been one of the principle genetic loci shaping human cultivation of barley over the last 10,000 years.

### Domestication of the six-row phenotype preserved VRS1 haplotype diversity in cultivated barley

All wild barley accessions analysed were phenotypically two-row and carried two-row *VRS1* haplotypes, indicating no instances of feral six-row individuals in our panel. Haplotype analysis landraces and cultivars identified all three previously described six-row alleles, *vrs1.a1, vrs1.a2* and *vrs1.a3*, consistent with previous studies of cultivated barley germplasm from this and other regions (Cuesta-Marcos et al. 2010; Komatsuda et al. 2007; Saisho et al. 2009; Sakuma et al. 2017). *VRS1* haplotype diversity declined sharply from wild barley to landraces (28 → 11), but only modestly from landrace to cultivars (11 → 7, with six-row haplotypes predominating in both of these genepools). This suggests that six-row mutations that arose post domestication were opportunistically selected due to their visually striking mutations, thus increasing haplotype diversity at the locus. Indeed, previous studies have suggested that the six-row alleles *vrs1.a2* and *vrs1.a3* originated via single point mutations from the corresponding wild-type *Vrs1.b2* and *Vrs1.b3* alleles in landraces from East Asia and the Western Mediterranean, respectively (Komatsuda et al. 2007). Our data broadly support these hypotheses: landraces predicted to carry the *vrs1.a2* allele originated from Spain, while haplotypes predictive of *vrs1.a3* occurred primarily in Russia and Eastern Europe (Figure 3).

### Evidence that the Vrs1.b1 allele arose in the wild barley genepool

The geographic origin of the *vrs1.a1* six-row allele has remained unclear (Komatsuda et al. 2007) due to the absence of a directly ancestral wild-type two-row haplotype representing the hypothetical *Vrs1.b1* allele. This is somewhat unexpected, given that *vrs1.a1* has been reported as the most common six-row allele in landraces (Saisho et al. 2009) and cultivars (Komatsuda et al. 2007). We also find *vrs1.a1* to be the most common landrace six-row allele, widely distributed across most of the geographic regions assessed here, suggesting it was the first to be domesticated. This is consistent with a recent genome-wide sequence-based estimate of barley haplotype origin around the *VRS1* locus which found the haplotype spanning *vrs1.a1* to be the most ancient of the three six-row alleles (Guo et al. 2025). Indeed, while only two six-row landraces within the Fertile Crescent were investigated here, both were found to contain haplotypes consistent with a *vrs1.a1* allele (Figure 3). However, we identified three wild barley accessions carrying a Vrs1.b1.1 haplotype, differing from the *vrs1.a1* by just the causative 1 bp exon-3 deletion that leads to a truncated VRS1 protein. Two of these Vrs1.a1.1 accessions were located in western Iran (Ilam and Khuzetan provinces) in the foothills of the Zagros mountains north-east of the Tigris river within the Fertile Crescent, while the third was from Turkmenistan near the Atrak river that forms the border between Turkmenistan and Iran, and situated to the south-east of the Caspian Sea. This Caspian Sea region also played a notable role in the domestication of the related temperate cereal crop, bread wheat (*Triticum aestivum*), which arose in Neolithic farmers’ fields via rare natural hybridization event(s) between cultivated tetraploid wheat and the wild diploid wheat *Aegilops tauschii* (Gaurav et al. 2022; Wang et al. 2013). Vrs1.b1.1 haplotypes were recently identified in two wild barley accessions via analysis of genomic sequence data generated uaing Illumina HiSeq 200 sequencing: FT897 and FT879 (ENA records ERX692648 and ERX692649, respectively). The collection sites for both were from the Karmanshah province of north-western Iran, within the foothills of the Zagros mountains. Collectively, these findings suggest that *Vrs1.b1* alleles were present at low frequency in stands of wild barley growing in regions of the Fertile Crescent where barley domestication occurred, as well as in the regions flanking the southern shores of the Caspian Sea where major Neolithic cereal domestication events occurred.

### Reduction and shift in two-row VRS1 haplotype diversity during domestication

In contrast to the emergence and proliferation of six-row alleles in the landraces, two-row *VRS1* haplotypes underwent a sharp reduction in diversity from wild barley to landraces (29 → 4), an expected outcome of domestication (e.g. Cheng et al. 2024; Civáň et al. 2024; Milner et al. 2019; Saisho et al. 2009). Two features stand out: Firstly, the predominant haplotype in wild barley (Vrs1.b2.2) sharply declined in frequency amongst landraces, and was completely absent in cultivars (although the *Vrs1.b2* allele was transmitted between the landrace and cultivar genepools via haplotype Vrs1.b2.1, albeit it at extremely low frequency in both germplasm groups). Secondly, the Vrs1.b3.1 two-row haplotype, absent from wild barley, predominates in both landrace (89%) and cultivar (94%) genepools. These patterns indicate that two-row haplotypes experienced a strong genetic bottleneck, likely driven by a founder effect followed by subsequent expansion of Vrs1.b3.1, potentially reinforced by human selection. Notably, haplotypes consistent with the *Vrs1.b2* allele carry two missense mutations (D8/G and D26/E) in the VRS1 N-terminus region also found in the rare *Vrs1.b1* allele, the *deficiens Vrs1.t1* allele, and the six-row *vrs1.a1* and *vrs1.a3* alleles that are characterized by truncated VRS1 proteins. Although these amino acid substitutions occur at relatively un-conserved residues within an unstructured region of the protein, they may be associated with pleiotropic effects. For example, the six-row mutant allele *vrs1.a1* (which shares both substitutions in the region upstream of the premature stop codon) is associated with reduced tillering, while the induced six-row mutants *vrs1*(*hex-v.3*) (a null *VRS1* deletion) and *vrs1*(*hex-v.6*) (derived from cv. Foma, *Vrs1.b3*) and so do not carry the D8/G and D26/E substitutions show no significant tiller reduction (Liller et al. 2015).

### VRS1 variation affecting protein sequence in wild barley

Ten wild barley haplotypes contained variants that altered VRS1 protein sequence. Allelic variation at *VRS1* is known to affect additional barley traits, making novel variants that affect VRS1 protein sequence of potential modern-day utility. Some of these pleiotropic traits are directly linked to the spikelets themselves, such as the extreme suppression of the infertile lateral spikelets observed in the *deficiens* allele *Vrs1.t1* (conferred by the S184/G missense mutation at a conserved amino acid position towards the C-terminus of the VRS1 protein) (Sakuma et al. 2017) and reduced lemma extension (Saisho et al. 2009). However, *VRS1* has also been shown to affect tiller number (Alqudah et al. 2016; Liller et al. 2015; Thirulogachandar et al. 2017), leaf size (Thirulogachandar et al. 2017) and plant height (Alqudah et al. 2016). Accordingly, *VRS1* negatively controls lateral floret development and leaf primordia initiation and development, where it is hypothesized to suppress the cell proliferation of leaf founder cells that form the leaf primordia. Four *VRS1* haplotypes carried missense mutations within the highly conserved homeobox-leucine zipper (HD-ZIP) domain, of which two (L73/V in Vrs1.b2.15 and D96/N in Vrs1.b2.20) fall in or near first two alpha helices known to play an essential role in protein positioning at the target DNA site in other HD-ZIP proteins. The proximity of the conservative L73/V substitution to the causal F75/L substitution underling the six-row *vrs1.a3* allele, and three independent six-row *VRS1* mutants (Komatsuda et al. 2007), underscores the domain’s functional importance. Additional wild barley mutations within the connecting loop between the second alpha helix and the recognition helix (non-conservative D96/N; haplotype Vrs1.b2.20) and close to the start of the recognition helix (conservative A101/T; haplotype Vrs1.b2.21) lie near residues where ≥16 induced missense mutants cause six-row or *intermedium* phenotype (see Supplementary Text 1) (Gottwald et al. 2009; Komatsuda et al. 2007). Substitutions at residue 107 exemplify this effect: non-conservative (R107/L from *hex-v.42* and *hex-v.43*) mutations confer six-row alleles, while conservative substitutions (R107/H from Barke TILLING line 11657-1) result in *intermedium* phenotype.

While the three haplotypes leading to amino acid changes in the N-terminus regions were not located within named protein domains, two are found within a region predicted to form an alpha fold structure at amino acid residues with high conservation across homologous plant proteins (conservative R31/K and non-conservative G43/R substitutions, originating from haplotypes Vrs1.b2.17 and Vrs1.b2.18, respectively). Finally, while the C-terminus region in which the three wild barley mutations affecting protein sequence (conservative A174/T, non-conservative G176/D, and conservative V178/I substitutions) does not contain notable 3D structure, much of this region is relatively highly conserved across VRS1 homologues. The importance of this region is highlighted by the presence of the non-conservative D184/G amino acid mutation underlying the *deficiens* allele *Vrs1.t1* (Gottwald et al. 2009; Hansson et al. 2024), as well as the intermedium mutation C194/S in *vrs1*(*Int-d.11*) (Komatsuda et al. 2007). Indeed, removal of the end of the recognition alpha-helix and downstream C-terminus region prevents interaction with the floral development gene *HvMADS13* and the inhibition of *HvMADS13* expression associated with lateral floret abortion (Shen et al. 2025). Collectively, the naturally occurring *VRS1* variants identified here in wild barley often cluster in conserved, functionally important regions and may influence key developmental traits, warranting further study of their phenotypic effects.

## Author contributions

JB undertook haplotype analyses and geographic plotting. HJ provided project resources. JC undertook sequence and bioinformatic analysis. The AEGIS Consortium facilitated discussions and provided the wider scientific context in which the work was analysed and interpreted. JC wrote the manuscript. JB, HJ and JC edited the manuscript.

## Acknowledgements

The work was supported via the Natural Environment Research Council via the project ‘The Domestication of Europe’ and by ‘AEGIS: Ancient Environmental Genomics Initiative for Sustainability’ (Novo Nordisk Foundation grant code NNF24SA0092560; Wellcome Trust grant code 313162/Z/24/Z). We thank the genebanks and institutes that provided barley germplasm: INRA (National Institute of Agricultural Research Biological Resource Centres, France), IPK (Leibniz Institute of Plant Genetics and Crop Plant Research genebank, Germany), IRTA (Institute of Agrifood Research and Technology, Spain), JHI (James Hutton Institute, UK), JIC GRU (John Innes Centre Germplasm Resource Unit, UK, a National Bioscience Research Infrastructure supported by the UKRI-BBSRC, grant number BBS/E/JI/23NB0001 for conserving and supplying germplasm through https://www.seedstor.ac.uk), NordGen (Nordic Genetic Resources Center, Sweden), RAC (Station de Recherche Agroscope Changins, Switzerland), USDA NSGC (U.S. Department of Agriculture National Small Grains Collection, USA), VIR (N.I. Vavilov Research Institute of Plant Industry genebank, Russia).

## Data availability

Genomic DNA sequences for all cultivars and one exemplar from each *VRS1* haplotype identified in landrace and wild barley accessions have been deposited in GenBank using the accession numbers listed in Supplementary Table S1. Germplasm is available via the relevant genebanks and institutes, as listed in Supplementary Table S1.

