## Supplementary Figure S1 for "The barley ear row-number allele ancestral to the six-row allele *vrs1.a1* is found in wild barley from the Fertile Crescent"

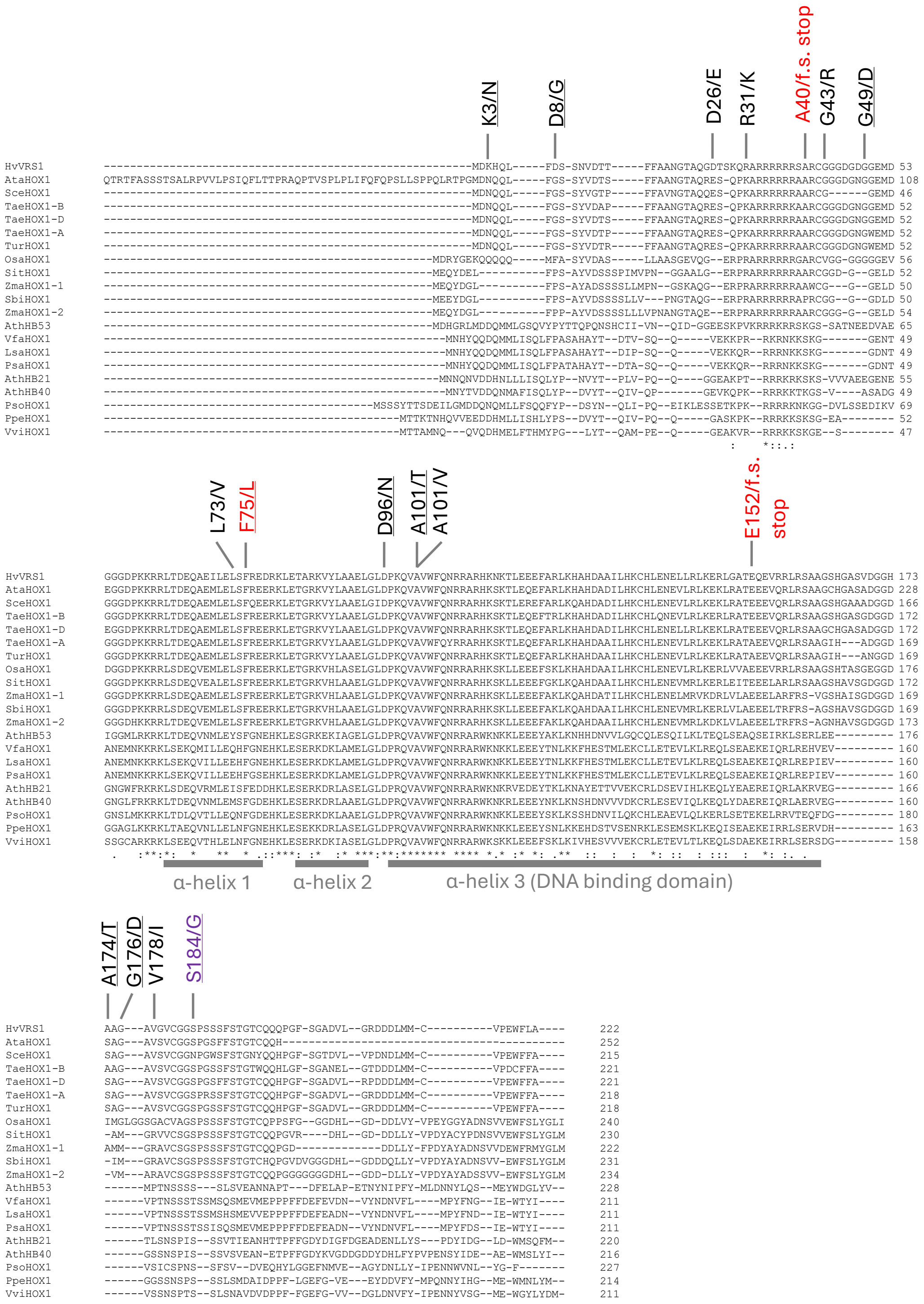


**Supplementary Figure S1.** Multiple protein alignment of plant VRS1 homologues. The positions of the predicted protein sequence variants identified are highlighted, as are the positions of the three alpha (α) helixes within the homeobox-leucine zipper domain (Panther ID PTHR24326). The causative mutations underlying the six-row *vrs1.a1* (E152/f.s. stop), *vrs1.a2* (A40/f.s. stop) and *vrs1.a3* (F75/L) are highlighted in red. The mutation underlying the two-row *deficiens* allele is highlighted in purple. Non-conservative amino acid substitutions are underlined. f.s. = frame shift. Stop = premature stop codon. Species abbreviations and protein accession numbers: HvVRS1 (*Hordeum vulgare*; BAH24152.1), AtaHOX1 (*Aegilops tauschii*; AET2Gv20863900.1), SceHOX1 (*Secale cereale*; SECCE2Rv1G0113870.1), TaeHOX1-B (*Triticum aestivum*; TraesCS2B02G405700.1), TaeHOX1-D (*Triticum aestivum*; TraesCS2D02G385500.2), TaeHOX1-A (*Triticum aestivum*; TraesCSU02G009800.2), TurHOX1 (*Triticum urartu*; TuG1812G0200004383.01.T01), OsaHOX1 (*Oryza sativa*; BGIOSGA026014-PA), SitHOX1 (*Setaria italica*; KQL26425), ZmaHOX1-1 (*Zea mays*; Zm00001eb107800_P001), SbiHOX1 (*Sorghum bicolor*; EER97443), ZmaHOX1-2 (*Zea mays*; Zm00001eb325630_P003), AthHB53 (*Arabidopsis thaliana*; AT5G66700.1), VfaHOX1 (*Vicia faba*; Vfaba.Hedin2.R1.1g361760.1), LsaHOX1 (*Lathyrus sativus*; CAK8539850.1), PsaHOX1 (*Pisum sativum*; Psat2g161280.1.cds), AthHB21 (*Arabidopsis thaliana*; AT2G18550.1), AthHB40 (*Arabidopsis thaliana*; AT4G36740.1), PsoHOX1 (*Papaver somniferum*; RZC55109), PpeHOX1 (*Prunus persica*; ONH96752), VviHOX1 (*Vitis vinifera*; Vitis04g01393.t01.CDS).
